# Shortwave Infrared Fluorescence Imaging with the Clinically Approved Near-Infrared Dye Indocyanine Green

**DOI:** 10.1101/100768

**Authors:** Jessica A. Carr, Daniel Franke, Justin R. Caram, Collin F. Perkinson, Vasileios Askoxylakis, Meenal Datta, Dai Fukumura, Rakesh K. Jain, Moungi G. Bawendi, Oliver T. Bruns

## Abstract

Fluorescence imaging is a method of real-time molecular tracking *in vivo* that has enabled many clinical technologies. Imaging in the shortwave infrared region (SWIR, 1-2 μm) promises higher contrast, sensitivity, and penetration depths compared to conventional visible and near-infrared (NIR) fluorescence imaging. However, adoption of SWIR imaging in clinical settings has been limited, due in part to the absence of FDA-approved fluorophores with peak emission in the SWIR. Here, we show that commercially available NIR dyes, including the FDA-approved contrast agent indocyanine green (ICG), exhibit optical properties suitable for *in vivo* SWIR fluorescence imaging. Despite the fact that their emission reaches a maximum in the NIR, these dyes can be imaged non-invasively *in vivo* in the SWIR spectral region, even beyond 1500 nm. We demonstrate real-time fluorescence angiography at wavelengths beyond 1300 nm using ICG at clinically relevant doses. Furthermore, we show tumor-targeted SWIR imaging with trastuzumab labeled with IRDye 800CW, a NIR dye currently being tested in multiple phase II clinical trials. Our findings suggest that high-contrast SWIR fluorescence imaging can be implemented alongside existing imaging modalities by switching the detection of conventional NIR fluorescence systems from silicon-based NIR cameras to emerging indium gallium arsenide (InGaAs) SWIR cameras. Using ICG in particular opens the possibility of translating SWIR fluorescence imaging to human clinical applications.

## Introduction

Fluorescence imaging in the near-infrared (NIR, 700-1000 nm) has enabled new technologies for preclinical and clinical applications.^1–5^ Compared with conventional diagnostic imaging such as computed tomography, positron emission tomography, or magnetic resonance imaging, NIR fluorescence imaging provides a lower cost, high sensitivity method for real-time molecular imaging.^2,3,5^ NIR fluorescence is also being evaluated in clinical trial for numerous image-guided surgery applications.^4,6,7^ A variety of NIR fluorophores are commercially available that exhibit high brightness, and the ability to target a range of biological substrates. Moreover, one of these dyes, indocyanine green (ICG), has been approved by the FDA for clinical use since 1959. ICG and other NIR dyes such as IRDye 800CW are the subject of over 300 clinical trials for applications such as fluorescence angiography and perfusion assessment in reconstructive and bypass surgeries, metastatic lymph node mapping and lymphatic transport in lymphedema, cancer localization and surgical margin assessment, and many others.^5–11^

Recent research has shown that extending fluorescence imaging into shortwave infrared wavelengths (SWIR, 1000-2000 nm) can further enhance the advantages of NIR imaging.^12–15^ Low levels of background tissue autofluorescence in the SWIR increase imaging sensitivity to a target fluorophore, and the different tissue absorption and scattering properties increase contrast of structures at greater penetration depths.^13,16–21^ However, the adoption of SWIR fluorescence imaging into clinical settings has been prevented by the limited availability of SWIR detection technology and the perceived need for FDA-approved fluorophores with peak emission in the SWIR spectral region. The availability of SWIR detectors, controlled in part by national defense-related policies such as the U.S. International Traffic in Arms Regulations (ITAR), is rapidly increasing, as over 20 SWIR cameras are now classified for dual (military and commercial) use by the U.S. Department of State.^22^ In addition, improvements in indium gallium arsenide (InGaAs) sensor fabrication, increasing supply, and growing markets have significantly decreased SWIR detection technology costs.^23^ As for SWIR-fluorescent probes, several examples of inorganic nanomaterials and hydrophobic organic molecules exist with peak emission in the SWIR. Increasing the quantum yield, functionality, and biocompatibility of SWIR fluorophores is an active focus of emerging research studies.^24–37^

Here we show that two commercially available dyes with peak emission in the NIR spectral region, including the FDA-approved contrast agent ICG, can function as SWIR emitters. Our findings are based on the observation that even though the emission spectrum of NIR dyes such as ICG or IRDye 800CW peaks outside of the SWIR spectral region, their fluorescence exhibits a broad shoulder with a spectral tail extending well into the SWIR that can be easily detected by modern SWIR cameras. By demonstrating both functional and targeted SWIR imaging, our findings suggest that NIR dyes could bridge the gap between current shortcomings of fluorescent SWIR probes and applications in clinical settings.

## Results and Discussion

### NIR dye emission detected with InGaAs

The conventional use of silicon-based technology to detect the emission spectrum of ICG has led many of approximately 3000 papers published on this dye to under-detect the extent of its SWIR emission. According to the Franck-Condon principle (or mirror image rule), most fluorescent organic dye emission spectra should approximate the mirror image of their absorption spectra.^38^ However, spectra of NIR dyes reported in the literature often appear to disobey this rule; while the absorption spectra of the dyes exhibit a clear shoulder to the blue of the absorption maximum, the reported emission spectra lack this shoulder. This disparity can originate from recording the emission spectra of NIR dyes on a silicon-based detector. The detection efficiency of these detectors sharply declines beyond 900 nm, limiting detection of the full tail of NIR dye emission.

Recording the emission spectrum on a more suitable detector for sensing SWIR emission, such as an InGaAs-based detector, shows that ICG emission indeed follows the Franck-Condon principle, and that the tail of its emission spectrum extends well into the SWIR region (Figure 1).^39^ Under diffuse 808 nm excitation, it is even possible to detect emission from an aqueous solution of ICG on an InGaAs SWIR camera beyond 1500 nm, even though the emission of ICG peaks at 820 nm. This finding is significant in the context of recent studies showing the improvement in contrast, sensitivity, and penetration that can be gained by performing fluorescence imaging at the longest wavelengths of the SWIR.^12–14^ The same principle applies not only to ICG, but also to other NIR dyes, such as IRDye 800CW (**Supplementary Figure 1**), which is currently in multiple phase II clinical trials and promises improved stability compared to ICG.^40^

**Figure 1.**
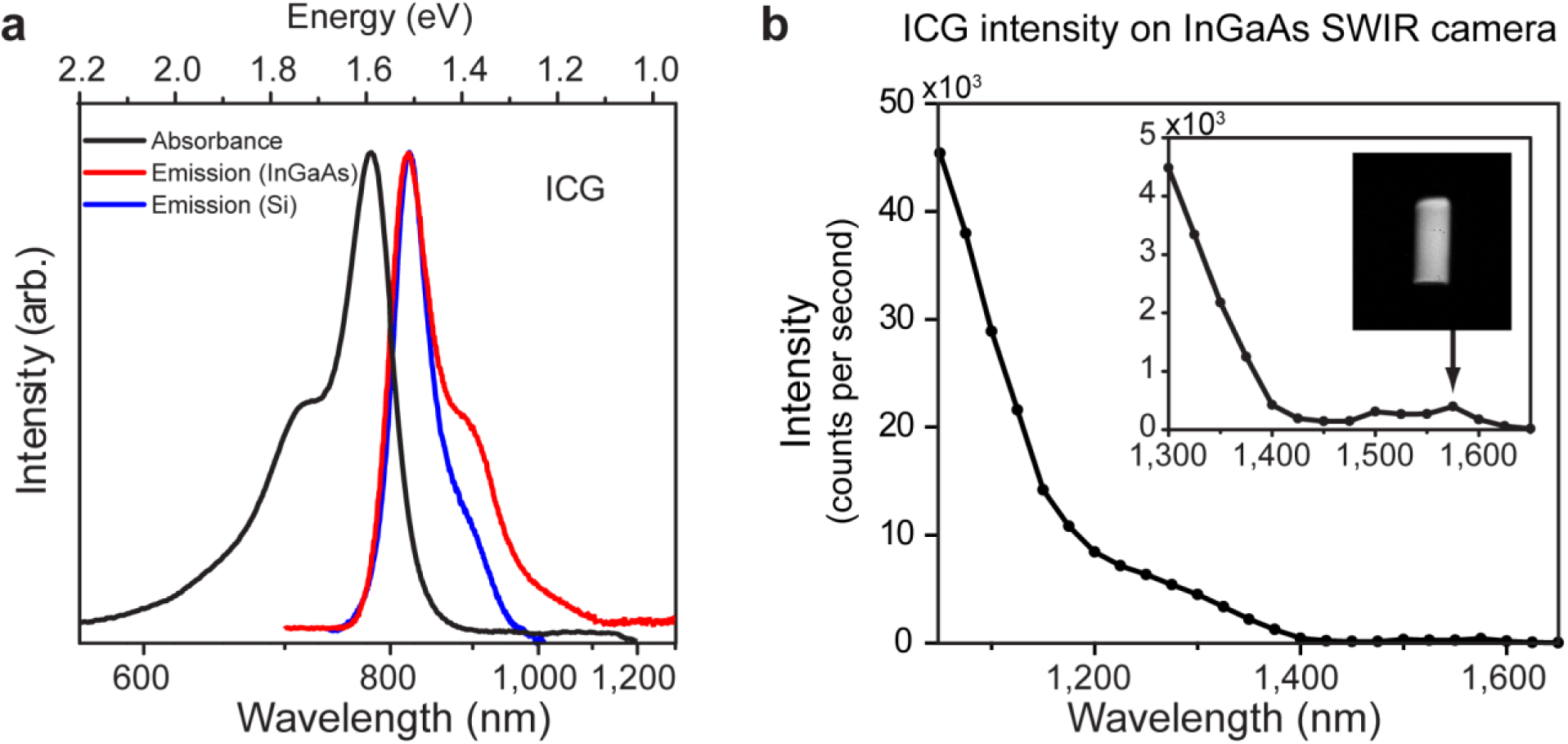
SWIR emission of the NIR dye indocyanine green (ICG). (a) Measuring the emission of ICG using a silicon-based detector (blue line) artificially truncates the low energy shoulder of the emission. Measuring the emission on an InGaAs detector (red line), which has greater sensitivity in the SWIR spectral region, recovers this emission tail, such that the full emission spectrum mirrors the absorption spectrum of ICG (black line), as predicted by the Franck-Condon principle. (b) The emission intensity of a 0.027 mg/mL aqueous ICG solution as detected in 20 nm spectral bands on an InGaAs camera, normalized by integration time, shows that emission from an aqueous solution of ICG is detectable up to 1575 nm (inset image of vial). The lower intensity between 1400 nm and 1500 nm is due to the absorption band of water in that region.

In fact, the SWIR emission from NIR dyes such as ICG and IRDye 800CW exceeds the brightness of a commercially available, state-of-the-art organic fluorophore developed specifically for *in vivo* SWIR imaging applications (IR-E1050) (Figure 2). We compare the optical properties of aqueous ICG and IRDye 800CW to IR-E1050, which has a peak emission in the SWIR.^41^ The brightness of a fluorescent probe is given by the product of the fluorescence quantum yield with the probe’s absorption cross-section at the excitation wavelength. ICG and IRDye 800CW exhibit both higher quantum yields in aqueous solutions (0.9% and 3.3% respectively), and higher peak absorption cross-sections (15 x10^4^ M^−1^ cm^−1^ and 24 x10^4^ M^−1^ cm^−1^) than IR-E1050 (quantum yield 0.2%, peak absorption cross section 0.80 × 10^4^ M^−1^ cm^−1^).^42–45^ As a consequence, when normalized to equimolar concentrations in water, the emission intensity of ICG between 1000 nm and 1300 nm is 8.7 times higher than for IR-E1050 (Figure 2). Using the absorption cross-section, quantum yield, and the ratio of the fluorescence signal between 1000 nm and 1300 nm to the total fluorescence signal (5% for ICG, 47% for IR-E1050) predicts that ICG should be 9.1 times brighter that IR-E1050, consistent with our measurements. We further compare the emission intensity of ICG, IRDye 800CW, and IR-E1050 in bovine blood on an InGaAs camera to best estimate the *in vivo* brightness of the probes (Figure 2). It is important that a full characterization includes a comparison in blood, as characterization in water underrepresents the *in vivo* brightness of ICG due to formation of H-dimer species at low concentrations in water that cause fluorescence quenching.^39^ To quantify the molar brightness of the probes on the camera, the intensity of each vial containing equal masses of the respective dye was measured individually, averaged, normalized to integration time, and multiplied with the molecular mass of the respective dye. Equimolar ICG and IRDye 800CW were 25 and 3.8 times brighter, respectively, than IR-E1050 in the wavelength range between 1000-1620 nm and 16 and 1.3 times brighter in the wavelength range between 1300-1620 nm when excited with 808 nm light.

**Figure 2.**
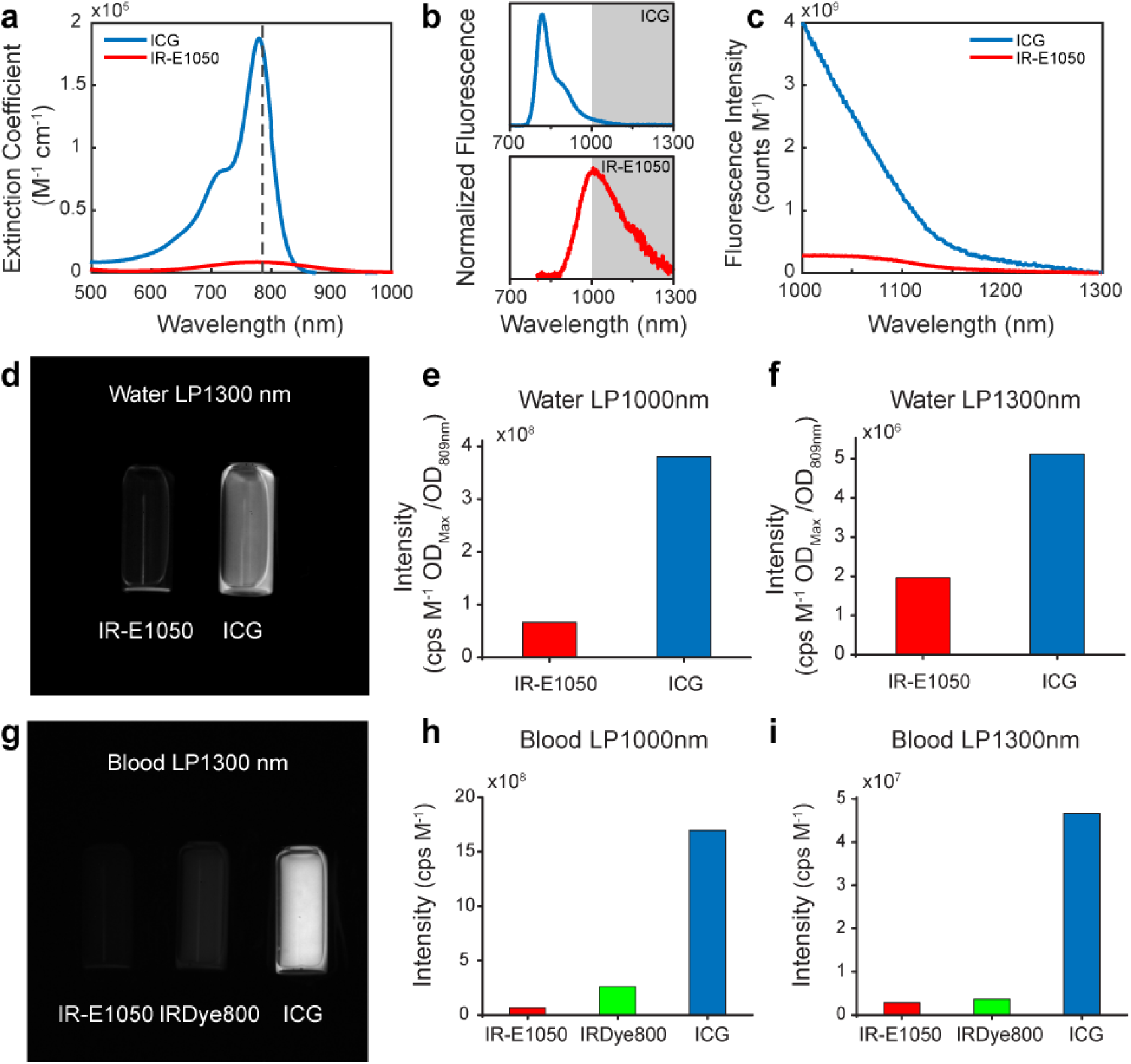
Comparison of the optical properties of ICG to other NIR and SWIR dyes. In addition to fluorescence quantum yield and spectral overlap with the SWIR region, SWIR brightness of a fluorophore also depends on absorption cross-section at the excitation wavelength. (a) Normalized to identical mass concentrations, ICG absorbs much more strongly than IR-E1050, a commercially available SWIR dye marketed for *in vivo* SWIR imaging applications, at 785 nm excitation (dashed line). In addition, ICG exhibits a higher fluorescence quantum yield (0.9%) than IR-E1050 (0.2%). (b) Whereas, the peak of the emission spectrum of ICG is significantly blue-shifted compared to IR-E1050, (c) the absolute measured emission intensity between 1000 nm and 1300 nm normalized to equimolar concentration is 8.7 times higher for ICG than for IR-E1050. (d) We further compared the fluorescence intensity of 0.01 mg/mL aqueous solutions of IR-E1050 and ICG on an InGaAs camera with 808 nm excitation, imaging at wavelengths between 1000-1620 nm. (e-f) The results also show superior emission of ICG compared to IR-E1050 both (e) between 1000-1620 nm (LP1000nm) and (f) between 1300-1620 nm (LP1300nm). (g) The high brightness of ICG in the SWIR is further increased in blood,^6^ so we compared the fluorescence intensity of IR-E1050, (0.01 mg/mL), with IRDye 800CW PEG (0.01 mg/mL, not accounting for PEG shell of 25-60 kDA), and ICG (0.01 mg/mL) in bovine blood. On a molar basis, the NIR dyes outperform IR-E1050 both beyond (h) 1000 nm and (i) 1300 nm. Standard deviations for signal intensity were found to be less than 5% for measuring signal intensity in blood and less than 15% for measurements in water.

Thus, commercially available and clinically-relevant NIR fluorophores have significant SWIR emission, eliminating one of the barriers to the adoption of SWIR fluorescence imaging in both research and clinical applications. These results further emphasize that the selection of contrast agents for SWIR fluorescence imaging is not limited to probes with peak emission in the SWIR. Due to the low quantum yields of conventional SWIR fluorophores, it is possible for the brightest probe to be one with only tail emission in the SWIR. Any new SWIR-fluorescent contrast agent should therefore be benchmarked against the SWIR emission of ICG in blood, as it is sufficiently bright for *in vivo* applications, and it is already FDA-approved and clinically used.

### High contrast SWIR fluorescence imaging *in vivo* using ICG

The SWIR emission of commercially available, FDA-approved, and clinically-used ICG enables straightforward application to *in vivo* fluorescence imaging in the SWIR. We present here a selection of *in vivo* imaging applications in multiple mouse models using the clinically approved dose of ICG, and demonstrate advantages of imaging ICG using SWIR detection over conventional NIR detection.

We show that imaging ICG in the SWIR enables high-contrast mesoscopic imaging of brain and hind limb vasculature in mice through intact skin (Figure 3), as has been previously demonstrated with carbon nanotubes.^13^ For this, we injected an aqueous solution of ICG into the tail vein of mice at a dose of 0.2 mg/kg, which is within the recommended dose for humans (0.2-0.5 mg/kg recommended, 5 mg/kg maximum).^6^ We illuminated the mice with 50-70 mW/cm^2^ of 808 nm excitation light, staying below the maximum permissible exposure limit (330 mW/cm^2^ for 808 nm continuous wave light).^46^ We noninvasively imaged the resulting fluorescence on a silicon camera at NIR wavelengths, and on an InGaAs camera at SWIR wavelengths between 1300 nm and 1620 nm, the detection cutoff of the cooled SWIR camera. We chose the 1300-1620 nm wavelength range, since as we show here and others have previously demonstrated, contrast and resolution are maximized above 1300 nm.^13,19,47^ We quantified the contrast within a region of interest containing the brain vasculature in the NIR image and the SWIR image by calculating the coefficient of variation, defined as the standard deviation of pixel intensity normalized to the mean pixel intensity (**Supplementary Figure 2**). We find that the SWIR image contrast for brain vasculature is nearly 50% greater at a value of 0.29, compared to the NIR image with a contrast value of 0.20, and it is more than 58% greater for hind limb vasculature at a value of 0.19 for SWIR imaging compared to 0.12 for NIR imaging. Furthermore, we calculated the apparent vessel width for a brain vessel by measuring the full width at half maximum of a two-Gaussian fit to the intensity profile across a brain vessel of interest. We find that the apparent vessel width in this specific example is over twice as wide in the NIR image as in the SWIR image with values of 430 μm and 210 μm, respectively. These findings are in good agreement with previous studies employing other fluorophores.^13^ Thus, contrast and resolution of fine vasculature structures can be greatly improved while using FDA-approved ICG contrast by simply switching the detection wavelength from traditional NIR imaging using a silicon camera to detection beyond 1300 nm on an InGaAs SWIR camera.

**Figure 3.**
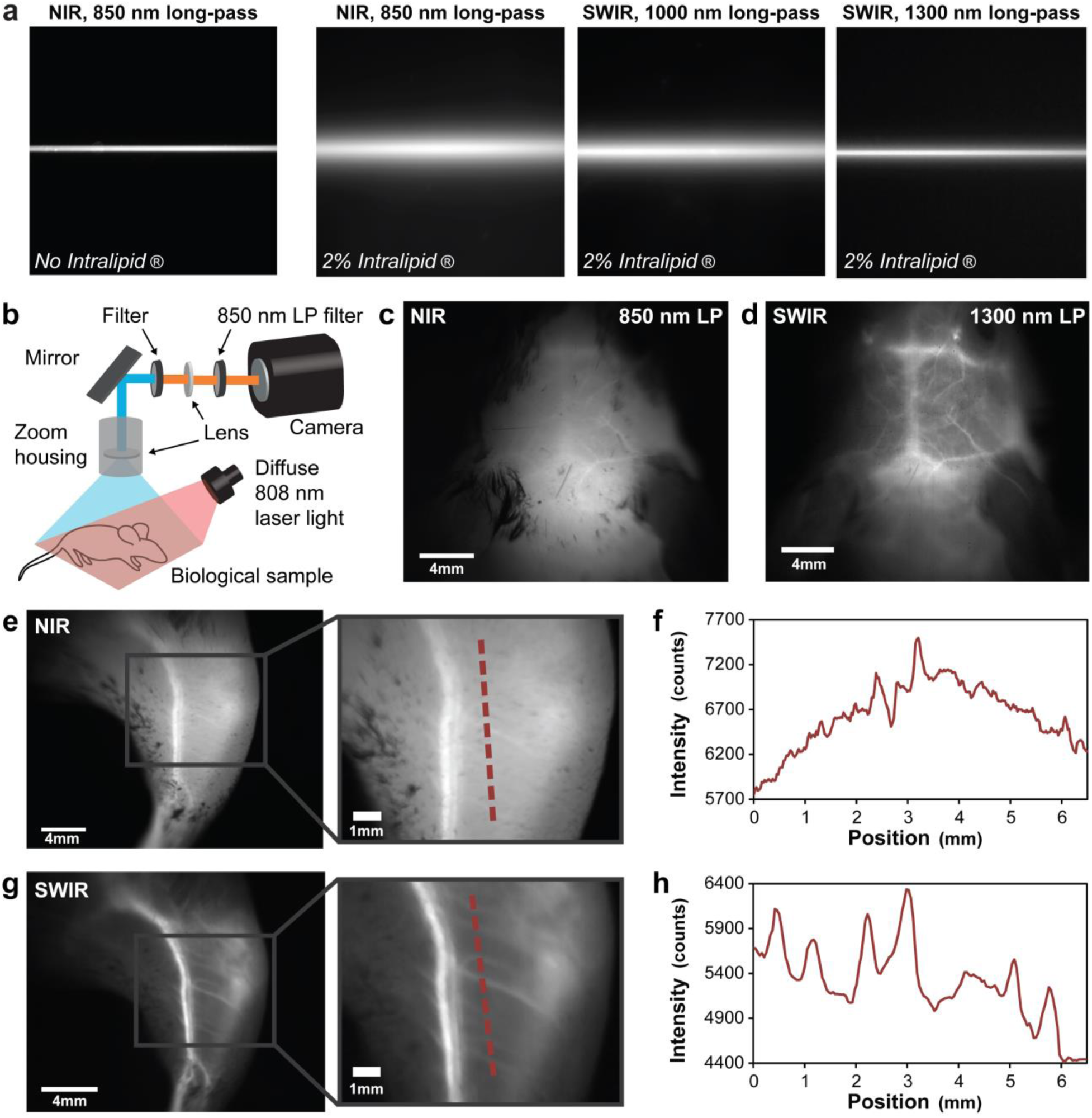
High contrast SWIR fluorescence imaging using ICG. (a) A 0.8-1.1 mm wide (inner diameter) capillary was filled with aqueous ICG, and the fluorescence was imaged using 850 nm longpass NIR detection. The capillary was then submerged in approximately 3 mm of 2% Intralipid® liquid tissue phantom and imaged again with 850 nm longpass NIR detection, 1000 nm longpass SWIR detection, and 1300 nm longpass SWIR detection, showing that the scatter of light caused by the tissue phantom is suppressed with increasing imaging wavelength. (b) We developed an ICG fluorescence imaging set-up for *in vivo* imaging which includes either a silicon or InGaAs camera, an 850 nm longpass excitation filter, a commercial SWIR lens or custom lens pair, an optional longpass or band-pass filter to set the imaging wavelengths, and a mirror facing the imaging stage where the biological sample is illuminated with diffuse 808 nm light at 50-70 mW/cm^2^. (c) We imaged the brain vasculature of a mouse using ICG contrast and find that the vessels are difficult to resolve through skin and skull using NIR detection on a silicon camera. (d) Switching to 1300 nm longpass SWIR detection on an InGaAs camera greatly improves vessel contrast. (e) Similarly, only large hind limb vessels are imaged with good contrast through the skin using NIR detection. (f) The intensity across a line of interest shows insufficient contrast to resolve smaller vessels from background signal. (g) Using 1300 nm longpass SWIR detection greatly improves image contrast and (h) resolution of vessels. All images were scaled to the maximum displayable intensities.

We further show that the contrast improvement of SWIR detection over NIR detection can be enabling for microscopic imaging (**Supplementary Figure 3**). We incorporated ICG into polyethylene glycol-phosopholipids^48^ to increase the blood half-life, which is typically limited to 3-4 min.^49,50^ We injected an aqueous solution of these micelles into the tail vein of a mouse with an implanted cranial window and used a microscope to image the fluorescence of the ICG phospholipid micelles in the brain vasculature with both NIR and 1300 nm longpass SWIR detection. Images of the entire cranial window at two times magnification show the ability to resolve nearly all the same vessels using either NIR or SWIR imaging. The overall contrast however, was 1.4 times greater for the SWIR image (standard deviation/mean was 0.24 for NIR versus 0.33 for SWIR). Higher magnification (6 times) reveals that this contrast improvement using SWIR imaging enables the resolution of vessels which, due to the high label density of surrounding vessels, are difficult to distinguish from background signal in the NIR image.

These findings could improve clinical use of ICG in fluorescence angiography. Clinical imaging of ICG in the NIR has already shown value for angiography in ophthalmology, intraoperative assessment of blood vessel patency in tissue graft, bypass surgeries, and intracranial aneurism surgeries.^1,5,51–54^ Noninvasive imaging of lymphatic vasculature using ICG has also been described for surgical mapping and intraoperative identification of sentinel nodes, evaluation and monitoring of lymphedema, and following surgical excision of nodes.^1,5,55,56^ Increasingly, ICG fluorescence imaging is also being used in robot-assisted surgery, in which the surgeon relies on visual cues instead of tactile feedback.^7,11,56^ However, full clinical implementation has been partially limited by insufficient image quality in deep operating fields. The higher contrast of ICG SWIR fluorescence imaging over NIR imaging could benefit these applications and enable the resolution of finer vessels, especially in systems with high label density or interfering background signal. Importantly, the implementation of this contrast improvement would be straightforward, requiring only a switch from cameras with NIR detection to those with SWIR detection while continuing the familiar surgical set-up and use of ICG.

### Real-time SWIR fluorescence imaging using ICG

An essential component of fluorescence-guided surgery is the ability to perform real-time imaging. We show that the SWIR emission of ICG is sufficiently bright for real-time imaging at high frame rates using clinically approved doses of ICG. In one example, we performed heart angiography in mice. Using diffuse 808 nm excitation and a 1300 nm longpass emission filter, we were able to image the vasculature of the beating heart at speeds of 9.17 frames per second (**Supplementary Video 1**) while resolving fine vessels on the exterior surface against the underlying contrast (**Supplementary Figure 4**). The acquisition speed was sufficiently fast to capture the anesthetized mouse heart rate of 207 beats per minute, determined by tracking the intensity fluctuations of the heart (Figure 4a-c). Furthermore, the metabolism of the ICG was tracked as it reached the heart, lungs, peripheral veins, and was finally cleared by the liver (Figure 4d-g).

**Figure 4.**
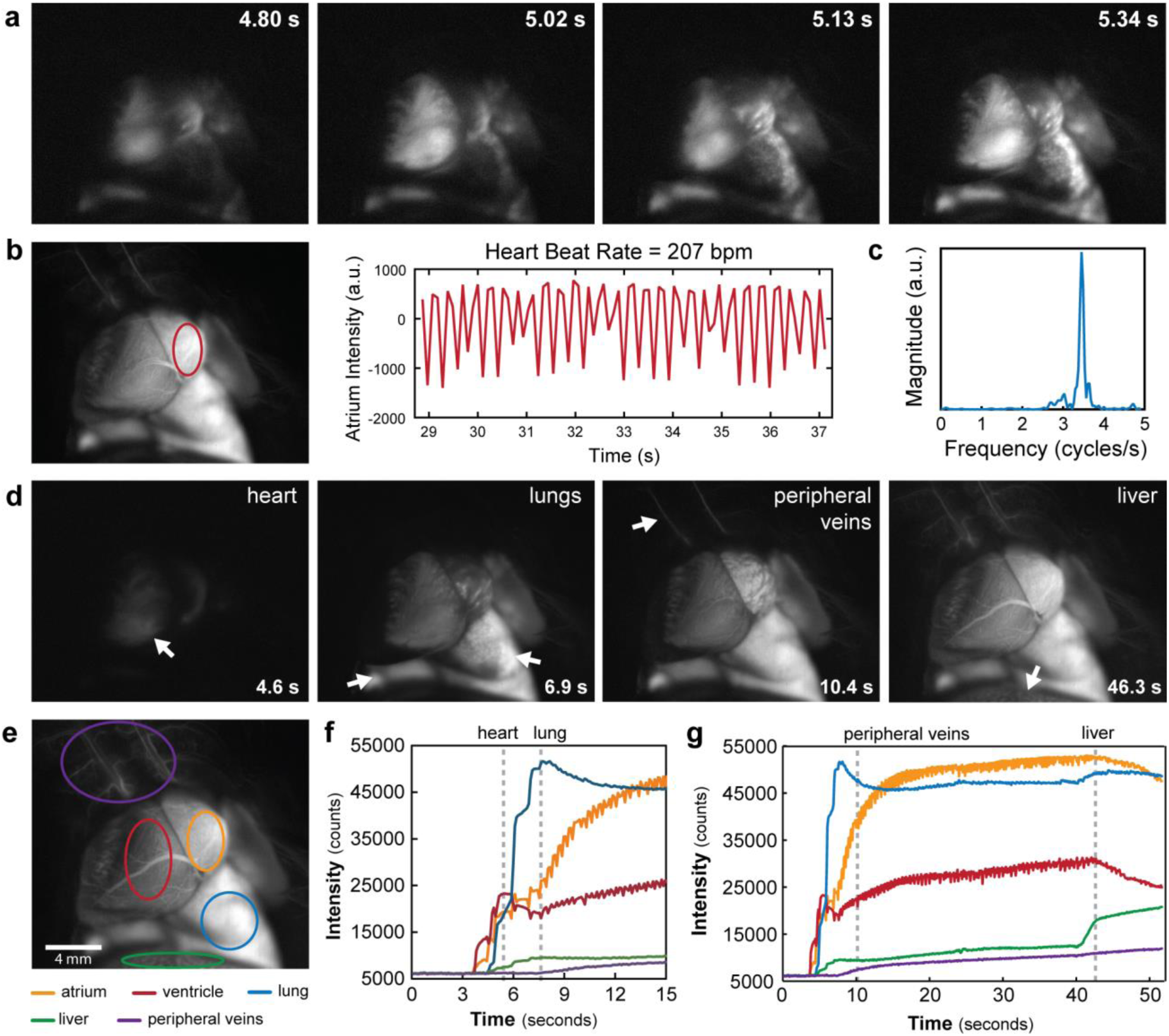
High temporal resolution demonstrated through ICG SWIR fluorescence angiography. SWIR fluorescence angiography was performed in a mouse heart at 9.17 frames per second using ICG for contrast, diffuse 808 nm excitation, and a 1300 nm longpass emission filter on an InGaAs SWIR camera (see **Supplementary Video 1**). (a) Temporal resolution was sufficiently high to resolve the heartbeat of the mouse. (b) By tracking the intensity fluctuations of the atrium of the heart, and (c) taking the Fourier transform, the heart rate was determined to be 207 beats per minute for the anesthetized mouse. (d) The ICG metabolism could also be tracked, as fluorescence first appears in the heart of the mouse, followed by the lungs, the peripheral veins, and finally the liver, which is the metabolic clearance method of ICG. (e) The fluorescence intensity of the heart atrium (orange), heart ventricle (red), lung (blue), liver (green), and peripheral vasculature (purple) are plotted to show the metabolism of the injection as it (f) arrives first at the heart and the lungs, and (g) at the peripheral vasculature before beginning to be cleared through the liver.

In a second example, we noninvasively image the hepatobiliary excretion of ICG into the small intestine.^6^ ICG emitted sufficient signal for near video rate imaging (19.7 frames per second, 1200 nm longpass filter), enabling capture of the peristaltic movements of the small intestine (**Supplementary Video 2**). Although at the cost of speed (2.0 frames per second), it was even feasible to image the ICG clearance using a 1500 nm longpass filter, capturing wavelengths between roughly 1500 and 1620 nm (**Supplementary Figure 5**).

In a third example, we noninvasively image lymphatic flow in a mouse. We injected an aqueous solution of ICG subcutaneously in the hind paws and tail, then image through intact skin the flow of lymphatic clearance (**Supplementary Video 3**). Using various bandpass filters across the SWIR, we find that lymph vessels and nodes are visible with ICG contrast up to approximately 1400 nm, at which point only the vessels and superficial nodes are visible and the signal of deeper lymph nodes becomes attenuated (**Supplementary Figure 6**). Thus, for applications which are not contrast-limited, such as those with low background signal and/or high label specificity, it may be preferable to image ICG in the NIR or the shorter wavelengths of the SWIR where signal attenuation is minimized.

These examples demonstrate the applicability of ICG SWIR imaging to fluorescence guided surgery. ICG is bright enough to image in the SWIR at speeds sufficiently high for intraoperative imaging of dynamic or moving features, shown here in the heart, small intestine, and lymphatic system of a mouse. We show that the required speed for a given application can be balanced with the desired contrast by selecting the imaging wavelength; using the full SWIR regime enables the highest frame rates due to maximized signal from ICG, while imaging at the longest SWIR wavelengths can be used to improve contrast at the cost of speed. Thus, using a SWIR camera for detecting ICG provides a tunable platform for optimizing both contrast and speed in fluorescence guided surgery.

### Targeted SWIR fluorescence imaging *in vivo* using IRDye 800CW

The ease of molecular targeting is one of the major strengths of fluorescence imaging over other imaging modalities. We show here that straightforward conjugation chemistry can be used to perform targeted imaging in the SWIR using the NIR dye IRDye 800CW. We used a commercially-available labeling kit to conjugate IRDye 800CW to the tumor-targeting antibody trastuzumab.^57^ We injected the dye-antibody conjugate into mouse models implanted with human BT474 breast cancer cells in the brain and noninvasively imaged the SWIR fluorescence emitted from the IRDye 800CW-labeled tumor. Subsequently, we injected IRDye 800CW conjugated to polyethylene glycol (PEG), highlighting brain vasculature surrounding the tumor. A multi-color functional image of the brain was then generated by temporally resolving the two labels, i.e. by assigning different colors before and after the addition of IRDye 800CW PEG (Figure 5).^58^

**Figure 5.**
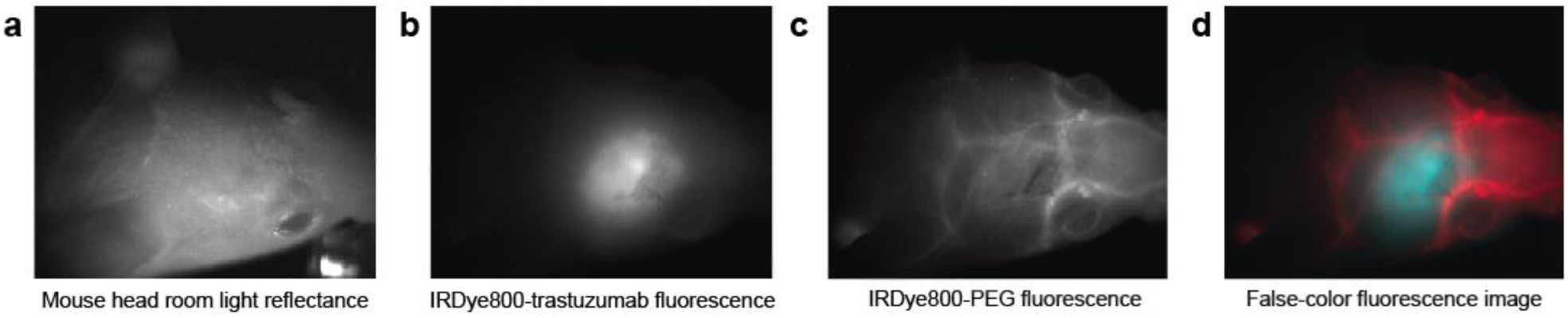
Targeted SWIR imaging *in vivo* with IRDye 800CW. (a) A nude mouse with a brain tumor from implanted human BT474 breast cancer cells is imaged here under room light illumination (bright field imaging). (b) Three days after injecting IRDye 800CW-trastuzumab conjugate, fluorescence from the labeled tumor was imaged on a SWIR camera with 1150 nm longpass emission detection. (c) Immediately subsequent injection of IRDye 800CW conjugated to polyethylene glycol (PEG) was used to provide contrast to surrounding brain vasculature which was temporally separated from the labeled tumor in post-processing. (d) The tumor and vessel images were then assigned a color (blue and red respectively) and overlaid to generate a false-color fluorescence image of the tumor and its surrounding vasculature.

While targeted SWIR imaging has previously been hindered by either the challenging preparation of targeted nanomaterials, size-dependent delivery effects, or the unavailability of commercial solutions, the use of small NIR dye labelling kits overcomes these barriers. NIR imaging of these readily available, targeted dyes has already shown promise for aiding cancer localization and intraoperative surgical margin assessment. Our results suggest that SWIR imaging of these NIR dyes could further benefit these applications by increasing the resolution of both fine and large structures which may be overlapping, as this is a common occurrence in highly vascular malignant lesions.

## Outlook

We show that established, commercially-available NIR dyes, including the FDA-approved dye ICG, can be used to perform state-of-the art SWIR imaging including noninvasive, real-time imaging in blood and lymph vessels, and molecularly-targeted *in vivo* imaging. The advantages of SWIR imaging over NIR techniques, such as increased sensitivity, contrast, and resolution of fine anatomical structures are therefore more readily available for increased adoption in pre-clinical and clinical imaging systems, simply by switching the detection from conventional silicon-based NIR cameras to emerging, high-performance InGaAs SWIR cameras. While no FDA-approved fluorophores with peak emission in the SWIR yet exist, we show here that detecting the off-peak fluorescence of clinically accessible NIR dyes on SWIR detectors bears the potential for rapid translation of SWIR fluorescence imaging to humans in clinical applications.

## Materials and Methods

### Absorption and emission characterization

An aqueous solution of ICG (Pfaltz & Bauer), IRDye 800CW PEG (LI-COR Biosciences), or IR-E1050 (Nirmidas Biotech) was prepared in SWIR-transparent quartz cuvettes. Absorption spectra were recorded on a Cary 5000 UV-Vis-NIR infrared spectrometer (Varian). To collect the emission spectra, samples were excited with a 635 nm diode laser (Thorlabs), and emission was collected using a pair of off-axis parabolic mirrors, and directed to a single-grating spectrometer (Acton, Spectra Pro 300i). An InGaAs calibrated photodiode (Thorlabs, DET10N) was used to detect the intensity of the emission. The tail emission of ICG was also detected using an InGaAs linear array (Princeton Instruments OMA-V). The emission spectrum of ICG was also measured using excitation from a 532 nm diode laser, and the resulting emission detected on a silicon-based spectrometer (OceanOptics, QE65000).

The emission intensity of ICG in water (0.027 mg/mL) was also measured in 20 nm spectral bands across the SWIR using an InGaAs SWIR camera (Princeton Instruments NIRvana 640). A glass vial containing the ICG solution was illuminated with 50-70 mW/cm^2^ of light from a 10?W 808?nm laser (Opto Engine; MLL-N-808) coupled into a 910μm-core metal-cladded multimode fiber (Thorlabs; MHP910L02) and diffused through a ground-glass plate (Thorlabs; DG10-220-MD). The peak emission of the laser was measured to be 809 nm. The emitted light was directed from the imaging stage to the camera using a four inch square first-surface silver mirror (Edmund Optics, Part No. 84448), then filtered through two colored glass 850 nm longpass filters (Thorlabs FGL850S) and through a liquid crystal tunable filter (PerkinElmer VariSpec LNIR, 20 nm bandwidth), and focused onto the SWIR camera with a C-mount objective (Navitar SWIR-35). The tunable filter was scanned from 900 to 1650 nm in 25 nm steps. A final image at 950 nm was acquired to ensure that the intensity matched the first measurement, verifying that the ICG solution was not bleached throughout the measurement.

### Photoluminescence quantum yield

Quantum yield measurements were obtained using an integrating sphere (Labsphere RTC-060-SF). The sample was illuminated using a 785 nm diode laser with an excitation power of 25 mW that was modulated at 260 Hz. The output was collected using a calibrated germanium detector (Newport: 818-IR) through a Stanford Research Systems lock-in amplifying system. Colored glass longpass filters (2x Schott Glass RG800 or 1x RG850) were used to block the excitation beam. The sample was placed in a PTFE capped quartz cuvette and a solvent blank was used to ensure a consistent environment inside the integrating sphere. The measured photocurrent was adjusted to account for the external quantum efficiency of the detector when calculating the quantum yield. Finally, the measured quantum yield was corrected to account for leakage of the excitation light and the transmittance of the filter.

While measuring the absolute quantum yield of IR-E1050 using an integrating sphere, we found a lower value (0.2%) compared to the one reported by the manufacturer (2%), which used the dye IR-26 as reference for relative quantum yield measurements. Their relative quantum yield measurements rely on a quantum yield of 0.5% for IR-26 that was reported by Drexhage and coworkers in 1982.^59^ More recently however, two independent publications by Beard and coworkers^60^ (0.05%) and Resch-Genger and coworkers^61^ (0.07%) have found the original value to be ∼ten times too large. Measuring the absolute quantum yield of IR-26 in our laboratory yielded a value of 0.05%, which is in line with the more recent publications. We therefore believe that 0.05% is the true value of the quantum yield of IR-26.

### Brightness comparison of ICG, IRDye 800CW, and IR-E1050

To compare the brightness in aqueous solutions, ICG (Pfaltz & Bauer) and IR-E1050 (Nirmidas Biotech) were diluted to a concentration of 0.01 mg/mL in cell culture grade water and in 1x PBS, respectively. To compare brightness in blood, ICG, IR-E1050, and IRDye 800CW PEG (LI-COR Biosciences) were diluted in bovine blood (Rockland Immunochemicals, sodium citrate-conjugated) to a concentration of 0.01 mg/mL. The solutions were imaged individually, and side-by-side, in the above-described set-up with 808 nm excitation, and the emission filtered through either two colored glass 1000 nm longpass filters or through an additional dielectric 1300 nm longpass filter. In the resulting background-corrected images, the average emission intensity was calculated from a region of interest within the vials and normalized to the integration time.

### In vivo imaging with ICG

All animal experiments were conducted in accordance with approved institutional protocols of the MIT Committee on Animal Care. No blinding or randomization was required for the animal studies. The *in vivo* imaging set-up is as described above with diffuse 808 nm excitation (50-70 mW/cm^2^), and an exchangeable filter holder for incorporating various bandpass and longpass emission filters. Different objectives were used depending on the requirements of each application. The InGaAs camera (Princeton Instruments, NIRvana 640) was cooled to −80 ^o^C, the analog to digital (AD) conversion rate set to 2 MHz or 10 MHz, the gain set to high, and different exposure times used to achieve sufficient signal and/or frame rates (**Supplementary Table 1**). All images were background-and blemish-corrected within the LightField imaging software. The silicon camera (Princeton Instruments PIXIS 1024BR) was cooled to −70°C, the AD conversion rate set to 2 MHz, the gain set to high, and the exposure time adjusted to achieve sufficient signal and/or frame rates (**Supplementary Table 1**). ImageJ was used to average 10 frames for NIRvana and PIXIS images. Frame averaging was not used in videos unless otherwise noted. ImageJ was also used for all image measurements (pixel intensity average, standard deviation, etc.).

Tissue phantom images were taken by submerging a capillary filled with 0.0175 mg/mL aqueous ICG (Pfaltz & Baur) in 20% Intralipid® (Baxter Healthcare Corporation, Deerfield, IL, USA) that was diluted to 2% Intralipid® in water. Images were taken in the NIR and the SWIR of the capillary before and after being submerged 3 mm below the surface of the 2% Intralipid® solution.

Noninvasive brain vasculature imaging was performed in two C3H/HeJ mice (21.7 g and 20.5 g, female, 10 weeks old, The Jackson Laboratory). Mice were anaesthetized, and the hair on the head was removed. Mice were placed in the imaging set-up, injected via the tail vein with up to 200 μL of a 0.025 mg/mL aqueous solution of ICG, and imaged using a pair of achromatic doublet lenses (Thorlabs, 75 mm and 150 mm EFL, 25.4 mm, C-coated), a hard-coated 850 nm dielectric longpass filter and either a second 850 nm longpass filter with the PIXIS silicon detector or a 1000 nm or 1300 nm longpass filter with the NIRvana InGaAs detector. Gaussian-fitting was performed in MATLAB using the Curve Fitting toolbox.

Intravital brain imaging was performed in a NU/NU nude mouse (female, 15 weeks old) with implanted cranial window (BT474-GFP-Glue implant). ICG-phospholipids were prepared by dissolving 4.8 mg ICG in a 2:1 mixture of chloroform and methanol, agitating with 23 mg polyethylene glycol-2000 phospholipids in chloroform (1,2-dioleoyl-sn-glycero-3-phosphoethanolamine-N-[methoxy(polyethylene glycol)-2000] (ammonium salt), Avanti Polar Lipids, #880130C), drying off the solvent under nitrogen flow, and resuspending the dried phospholipids in cell grade water. The aqueous ICG-phospholipid mixture was passed through a 0.22 μm syringe filter before injecting via the tail vein into the anaesthetized mouse. We used a Nikon Ti-E inverted microscope equipped with a Stage UP Kit (Nikon) and a backport adaptor to which an 808 nm laser diode coupled to a fiber was attached. To eliminate laser speckle we used an Optotune speckle remover (Edmund Optics; 88-397). We used a dichroic longpass filter (Thorlabs, DMLP900R for SWIR; Edmund Optics 69-906 for NIR) to direct the excitation light to the sample and a 850 nm longpass, 1000 nm longpass, and/or 1300 nm longpass filter (Thorlabs; FELH1000, FELH0850) to select the emission light. Imaging was done with a 2X (Nikon; CFI Plan Apo Lambda) or a 4X (Nikon; CFI Plan Apo Lambda,) objective with additional 1.5X magnification from the microscope.

Heart angiography was performed in three Friend Virus B NIH Jackson (FVB/NJ) mice (male, 18 weeks old, The Jackson Laboratory). Mice were anaesthetized and surgery performed to expose the heart. One or two 200 μL boluses of 0.25 mg/mL ICG in water were injected via the tail vein while imaging the emission. The objective used was composed of a pair of 200 mm focal-length biconvex lenses (Thorlabs, 25.4 mm, C-coated) and three translatable 500 mm focal-length biconvex lenses (Thorlabs, 50.8 mm, C-coated) to set the focus. Emission was filtered through a hard-coated 850 nm dielectric longpass filter and either a 1000 nm, 1150 nm, 1200 nm, or 1300 nm longpass filter (Thorlabs).

For ICG hepatobiliary clearance imaging, a NCr nude mouse (male, 14 weeks old, Taconic Biosciences Inc.) was anaesthetized using a ketamine/xylazine cocktail, placed in the imaging setup with a pair of achromatic doublet lenses (Thorlabs, 75 mm and 500 mm EFL, 25.4 mm, C-coated), and injected via a tail vein catheter with 200 μL of ICG at a concentration of 0.25 mg/mL in water. Hepatobiliary clearance of the ICG was then monitored and imaged over approximately 2 hours using an 850 nm dielectric longpass filter and a silicon detector, or an additional 1200 nm longpass or 1500 nm longpass filter with InGaAs detection.

Lymph node imaging was performed in a C57BL/6 mouse (male, 15 weeks old, The Jackson Laboratory). The mouse was anaesthetized and the hair was removed, and approximately 10 μL of 0.18 mg/mL aqueous solution of ICG was injected subcutaneously in the left and right hind footpad and tail of the mouse. The fluorescence was imaged with a commercial Navitar lens with 850 nm, 1000 nm, 1200 nm, 1300 nm, 1400 nm longpass dielectric filters (Thorlabs).

### In vivo targeted imaging with IRDye 800CW

IRDye 800CW was conjugated to trastuzumab (Genentech)^57^ using a labeling kit (Li-Cor, P/N 928- 38040). Briefly, 1 mg of trastuzumab was dissolved in 900 μL PBS buffer and 100 μL 1M potassium phosphate buffer (pH 9) to obtain a solution with a pH of 8.5. Twenty-five microliters of ultra-pure water were added to the NHS-IRDye 800CW and the solution was shaken for 2 minutes until the dye completely dissolved. Eight microliters of the resulting solution were added to the trastuzumab solution and the mixture was incubated for 2h at room temperature protected from light. Subsequently the trastuzumab-dye conjugate was separated from free residual dye using Pierce Zeba desalting spin columns.

Two tumor-bearing (BT474-Gluc) NU/NU nude mice^57^ (female, 11 weeks old) were injected intraperitoneally with 330 μL of IRDye 800CW-trastuzumab (15 mg/kg dose). Two additional tumor-bearing nude mice were not injected and served as controls. After three days, the mice were anaesthetized and imaged using diffuse 808 nm excitation, the previously described pair of achromatic doublet lenses, 850 nm and 1000 nm longpass dichroic filters, and an InGaAs camera. The mice were then injected via the tail vein with up to 300 μL of IRDye 800CW-PEG (Li-Cor, P/N 926-50401) and imaged with 850 nm longpass and either a 1000 nm, 1150 nm, 1200 nm, or 1300 nm longpass dichroic filter.

### Data availability

The authors declare that all data supporting the findings of this study are available within the paper, its Supplementary Information, and/or upon request from the authors.

## Acknowledgements

This work received support in part from the NIH (through the Laser Biomedical Research Center, Grant 9-P41-EB015871-26A1 to M.G.B.; CA080124 to R.K.J. and D.F.; CA126642 and CA197743 to R.K.J.; CA096915 to D.F.), the Massachusetts Institute of Technology through the Institute for Soldier Nanotechnologies, Grant W911NF-13-D-0001 (to M.G.B.), the NSF through EECS-1449291 (to M.G.B), the DOE Office of Science, Basic Energy Sciences, under Award No. DE-FG02-07ER46454 (to M.G.B., J.R.C.), and the DoD BCRP Innovator Award W81XWH-10-1-0016 (to R.K.J.). This work was further conducted with government support under and awarded by the Department of Defense, Air Force Office of Scientific Research, National Defense Science and Engineering Graduate Fellowship (NDSEG) 32 CFR 168a (to J.A.C.). D.F. was supported by a fellowship of the Boehringer Ingelheim Fonds. O.T.B. was supported by a European Molecular Biology Organization (EMBO) long-term fellowship. C.F.P. was supported by a NSF GRFP fellowship. M.D. was supported by the National Heart, Lung, and Blood Institute of the NIH (award number F31HL126449).

We would like to further thank Sylvie Roberge for excellent technical support and Tulio Valdez for helpful discussions on imaging applications. We would like to thank Ellen Sletten and Christopher Rowlands for critical feedback on this manuscript.

